# PyTorch Frustratometer: An Ultra-fast, Parallelized Tool for Proteome-Scale Local Energetic Frustration Analysis

**DOI:** 10.64898/2026.11.06.764892

**Authors:** Eli J. Draizen, Hana Jaafari, Carlos Bueno, Arnau Llaberia, Finley Clark, Ezequiel A. Galpern, Franco L. Simonetti, Maria I. Freiberger, Peter G. Wolynes, Diego U. Ferreiro, R. Gonzalo Parra

**Affiliations:** Evolutionary Systems Biophysics Group, Life Sciences Department, Barcelona Supercomputing Center, Barcelona, Spain; Computational Biology Group, Life Sciences Department, Barcelona Supercomputing Center, Barcelona, Spain; Center for Theoretical Biological Physics and Department of Chemistry, Rice University, Houston, TX, 77005, USA; Applied Physics Graduate Program, Smalley-Curl Institute, Rice University, Houston, TX, 77005, USA; Centre for Genomic Regulation, The Barcelona Institute of Science and Technology, 08003, Barcelona, Spain; Protein Physiology Laboratory, Departamento de Química Biológica - IQUIBICEN/CONICET, Facultad de Ciencias Exactas y Naturales, Universidad de Buenos Aires, C1428EGA, Buenos Aires, Argentina; Instituto de Química Biológica de la Facultad de Ciencias Exactas y Naturales, Consejo Nacional de Investigaciones Científicas y Técnicas - Universidad de Buenos Aires, C1428EGA, Buenos Aires, Argentina

## Abstract

Most natural protein sequences fold into well-defined three-dimensional structural ensembles, satisfying interactions that have been shaped over evolutionary timescales. The principle of minimal frustration states that most native interactions are energetically more favorable than alternative non-native ones, resulting in a funnel-shaped folding energy landscape. The FrustratometeR tool measures how the energetics of a residue or contact compares to decoys with different amino acid identities or configurations. Contacts are classified as minimally frustrated when these are more stable than random alternatives, highly frustrated when less stable, or neutral when energetically indistinguishable from them. Clusters of highly frustrated interactions are often evolutionarily conserved and are typically associated with functional regions. Therefore, identifying and quantifying local frustration can provide insights into both protein stability and function. Here we present a PyTorch reimplementation of FrustratometeR that accelerates conventional frustration calculations by ∼70–13,000× on a single GPU, depending on chosen frustration mode and protein size, with substantially larger gains for exhaustive mutational scanning, enabling proteome-scale local frustration analysis. We benchmark against the original algorithm on the CATH v4.4 S40 dataset and show near-identical results across all three frustration modes and mutational scanning. Source code is freely available at https://github.com/ParraLab-ESBG/pytorch-frustratometer.

## Introduction

For most proteins, molecular function is intimately linked to the adoption of well-defined three-dimensional structural ensembles. The principle of minimal frustration, a core concept of energy landscape theory, provides an explanation for why proteins can robustly fold into stable three-dimensional structures in biologically compatible times (Bryngelson and Wolynes, 1987). However, natural proteins are not completely frustration-free and retain *∼*10% of pairwise interactions in energetic conflict with their structural environment, i.e. they are frustrated (Ferreiro et al., 2007). Patches of these localized energetic conflicts are typically found at the surface of globules and are frequently associated with functional regions, including protein-protein interfaces (Ferreiro et al., 2007; Parra et al., 2015), allosteric sites (Ferreiro et al., 2011), catalytic residues (Freiberger et al., 2019), fuzzy interactions (Freiberger et al., 2021), and conformational switches (González-Higueras et al., 2024). These patches are also often evolutionarily conserved (Freiberger et al., 2023). Furthermore, local energetic frustration has emerged as a versatile approach for understanding diverse biological processes, including the molecular effects of disease-associated mutations (Kumar et al., 2016), the evolutionary relationship between exon organization and protein folding modularity (Galpern et al., 2024), the evolution and rewiring of protein binding specificity (Li et al., 2026b), and viral capsid assembly (Li et al., 2026a). More recently, frustration has also been exploited as a design principle to engineer protein dynamics and allosteric regulation (Broerman et al., 2025). Over the past two decades, local energetic frustration has evolved from a theoretical concept into a practical computational tool increasingly used in structural bioinformatics and computational biophysics. An overview of applications of local energetic frustration can be found in recent reviews (Parra et al., 2025; Parra and Ferreiro, 2025).

The original FrustratometeR introduced an accessible implementation of the AWSEM energy function (Davtyan et al., 2012) for estimating frustration directly from protein structures (Jenik et al., 2012), while subsequent developments expanded the methodology to incorporate electrostatic interactions, the analysis of molecular dynamics trajectories, and prediction of changes in frustration upon mutations analyses (Parra et al., 2016; Rausch et al., 2021).

Since the release of FrustratometeR, however, the scale of possible bioinformatic analysis in structural biology has changed dramatically. Advances in experimental structure determination continue to accelerate the growth of the Protein Data Bank, while AlphaFold2 (Jumper et al., 2021) and related methods have generated structural models for hundreds of millions of protein sequences (Lin et al., 2023; Varadi et al., 2022). At the same time, molecular dynamics simulations routinely produce conformational ensembles containing thousands of structures, and modern protein design approaches such as ProteinMPNN and diffusion-based generative models generate large libraries of candidate proteins for downstream evaluation (Dauparas et al., 2022; Watson et al., 2023). As a result, structural biology has shifted from analysing individual protein structures to interrogating datasets containing thousands or even millions of structures. In consequence, many analyses that were originally developed for individual proteins have become bottlenecks when applied at larger scales. This is particularly true for local energetic frustration, where estimating the energetic preference of residues or residue pairs requires evaluating large numbers of decoy states (Davtyan et al., 2012). Existing implementations of the algorithm are neither parallelized nor optimized for large-scale calculations, making them prohibitively expensive for proteome-scale analyses, large molecular dynamics trajectories, and protein design pipelines.

To address this challenge, we developed PyTorch Frustratometer, a complete reimplementation of the FrustratometeR framework designed for modern high-performance computing architectures. PyTorch Frustratometer preserves the established AWSEM energy model and frustration definitions while reformulating the underlying calculations as batched tensor operations executed natively on CPUs and GPUs. This redesign maintains numerical compatibility with previous implementations (Fig. 1G-I) while reducing computational cost by *∼*70–13,000*×*, depending on frustration mode and protein size (Fig. 3), enabling analyses that were previously computationally impractical.

**Figure 1.**
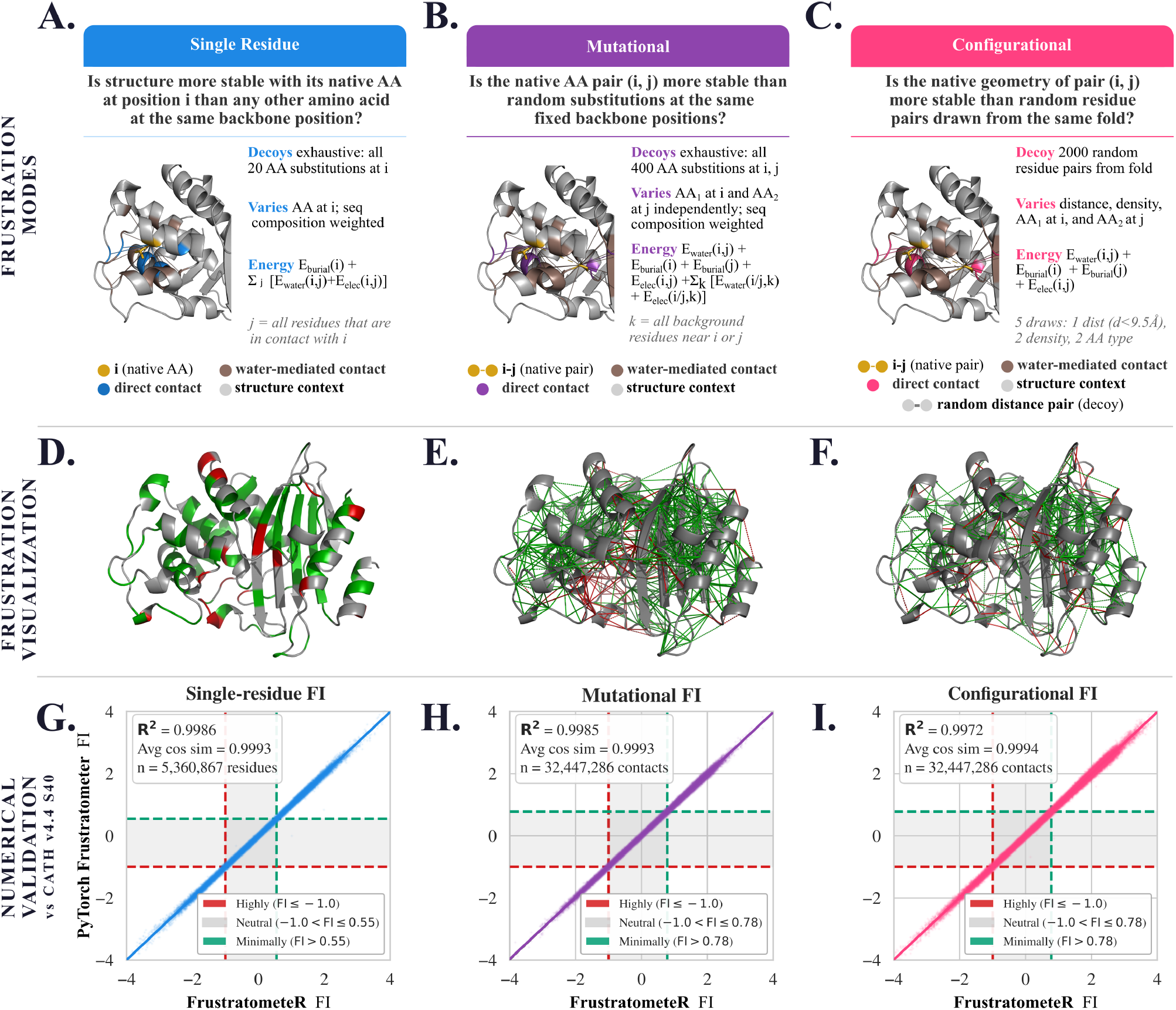
Frustration analysis modes and numerical validation of PyTorch Frustratometer. **(A–C)** Schematic representation of single-residue, mutational, and configurational frustration analyses, respectively, illustrating the native states, decoy ensembles, and energetic comparisons used to calculate each Frustration Index (FI), using the benchmark parameters of min-seq-sep=3, electrostatics=4.15, and burial-min-sep=3. Note that burial-min-sep was 3 by default in FrustratometeR, while it is now set to 1 (See methods section for details). Each colored line in the 3D representation is a contact used in the frustration calculation. **(D–F)** Corresponding frustration patterns mapped onto the structure of *Bacillus licheniformis β*-lactamase (PDB ID: 4BLM, chain A). For single-residue frustration (D), residues are colored according to their frustration class: red indicates highly frustrated residues, green minimally frustrated residues, and gray neutrally frustrated residues. For mutational (E) and configurational (F) frustration, lines connect interacting residue pairs, with red lines indicating highly frustrated contacts and green lines minimally frustrated contacts; neutral contacts are not shown. **(G–I)** Numerical comparison between PyTorch Frustratometer and FrustratometeR v1.4.1 across 34,653 CATH structures from CATH v4.4 S40 for single-residue, mutational, and configurational FIs, respectively. Dashed lines indicate the thresholds used to classify highly and minimally frustrated interactions.

Here we describe the implementation of PyTorch Frustratometer and evaluate its numerical agreement with FrustratometeR across all frustration modes, i.e. single-residue, configurational and mutational. We further show that GPU acceleration enables routine large-scale analyses, including exhaustive mutational scanning and proteome-scale frustration profiling, extending the scope of frustration analysis to massive datasets in modern structural bioinformatics.

## Results

### PyTorch Frustratometer re-implementation

PyTorch Frustratometer is a high-performance implementation of local energetic frustration analysis designed for proteome-scale applications. The python package preserves the energetic formulation and frustration indices of the original FrustratometeR while reformulating the calculations as vectorized tensor operations that can be efficiently executed on multicore CPUs and GPUs. Previous implementations accepted the input structures as PDB files only while the new PyTorch Frustratometer accepts input in mmCIF format as well. Output files follow the same format as in previous versions (ASCII), but we also include an option to write to Apache Parquet and Hierarchical Data Format, version 5 (HDF5; The HDF Group, 2026) when processing millions of proteins.

Protein structures and energy function parameters are represented as tensors, enabling the parallel evaluation of native and decoy interactions while eliminating most sequential computations. This implementation computes single-residue (Fig. 1A), mutational (Fig. 1B) and configurational (Fig. 1C) frustration indices (FIs) using the same definitions and classification criteria as previous versions, ensuring direct compatibility with existing analyses. Mutational and configurational frustration indices characterize the energetic frustration of pairwise residue contacts relative to alternative amino acid identities or interaction geometries, respectively. Because local frustration is fundamentally a contact-based property, its analysis at the level of contact maps is generally recommended, as highly frustrated interactions often cluster in specific regions associated with protein function; however, for simplicity and to facilitate comparisons with other residue-level biophysical features, a single-residue frustration index is also implemented, which has proven useful in different studies (Freiberger et al., 2019, 2023; Parra et al., 2025; Parra and Ferreiro, 2025). PyTorch Frustratometer outputs frustration values for all residues present in the protein structure (single-residue FI) as well as for each pairwise interaction (mutational and configurational FIs). As in previous versions, results can be visualized on top of the structures (Fig. 1D-F).

### Agreement with FrustratometeR

To verify that the new PyTorch Frustratometer produces results that are consistent with the FrustratometeR implementation, we compared frustration calculations across a diverse benchmark of 34,653 protein structures from CATH v4.4 S40 (Waman et al., 2025). We used identical input parameters for benchmarking: min-seq-sep=3, electrostatics=4.15, burial-min-sep=3 and for configurational mode, decoys=2000. Frustration indices computed by PyTorch Frustratometer showed almost perfect agreement with those obtained using FrustratometeR for contact level frustration (configurational R^2^ = 0.9972, see Fig. 1I; mutational R^2^ = 0.9985, see Fig. 1H), and single-residue (R^2^ = 0.9986, see Fig. 1G) frustration analyses. The small remaining differences are the consequence of the random sampling of decoy distributions. The actual values of FI are not expected to be exactly the same, as there is a random component in the sampling of the decoy states.

Frustration values were highly correlated between implementations, with 96% of residues assigned to the same frustration class (Supp. Fig. 1). Similarly, frustration values were highly consistent across independent runs despite the stochasticity introduced by random seed selection (Supp. Fig. 2). Minor numerical differences were limited to values close to the classification thresholds and arise from stochastic decoy sampling and floating-point arithmetic rather than methodological differences. These results show that PyTorch Frustratometer preserves the accuracy and interpretation of the original method while substantially improving computational performance. In addition to the common frustration modes, Guan et al. (2024) introduced the so-called smooth frustration mode, which is derived from the single-residue frustration index of two contacting residues that can be calculated as: F_smooth_= -(F_i_+F_j_) (see Methods, Supp. Fig. 3). This mode is now also implemented in PyTorch Frustratometer.

### Deep Mutational Frustration Scanning

The increased computational efficiency of PyTorch Frustratometer enables us to perform ultra-fast Deep Mutational Frustration Scanning (DMFS) analysis, named in analogy to Deep Mutational Scanning (DMS) experiments, in which changes in local frustration are calculated for every position and all alternative amino acid identities for a given structure. While this functionality is partially present in the previous version for user-selected positions, it can now be scaled to the entire protein at practically no additional computational cost. As an example, Fig. 2A shows the native single-residue frustration profile mapped onto the structure of *Bacillus licheniformis β*-lactamase (PDB ID: 4BLM, chain A). Fig. 2B shows the corresponding DMFS frustration matrix, highlighting regions surrounding the catalytic residues and the position for an experimentally determined WT–mutant structural pair for E166A in *Bacillus licheniformis* (see Supp. Fig. 4A–E for DMFS frustration matrices from different *β*-lactamases, including experimentally determined wild-type–mutant structural pairs). To assess the accuracy of this new functionality, we retrieved paired wild-type and mutant structures of experimentally resolved *β*-lactamases and compared their frustration values, explicitly calculated on those structures, with those estimated by DMFS from the corresponding wild-type structures (Supp. Table 1). Across all 25 WT–mutant pairs, DMFS-estimated and experimentally determined frustration values showed high agreement (R^2^ = 0.828; Fig. 2C, solid line correlation). One mutation deviated substantially from this relationship because it exhibited large structural rearrangements (D179Y) in the experimentally resolved mutant structure, changes that are known to be challenging to capture computationally (Kellogg et al., 2011) (Fig. 2C; Supp. Table 1). Thus, users should be aware that DMFS is limited in predicting changes in frustration when mutations substantially alter the local structural environment. Excluding this case resulted in an even higher agreement between DMFS estimates and the experimentally determined mutant structures (R^2^ = 0.969; n = 24; Fig. 2C, dashed line correlation), indicating that DMFS accurately captures changes in local frustration when mutations preserve the local structural environment. This new functionality allows users to rapidly estimate which amino acid substitution would lead to a desired change in frustration at a given position: e.g., at position 160, the WT amino acid is a neutral valine, with V160Y maximizing frustration and V160R minimizing it instead (Fig. 2B).

**Figure 2.**
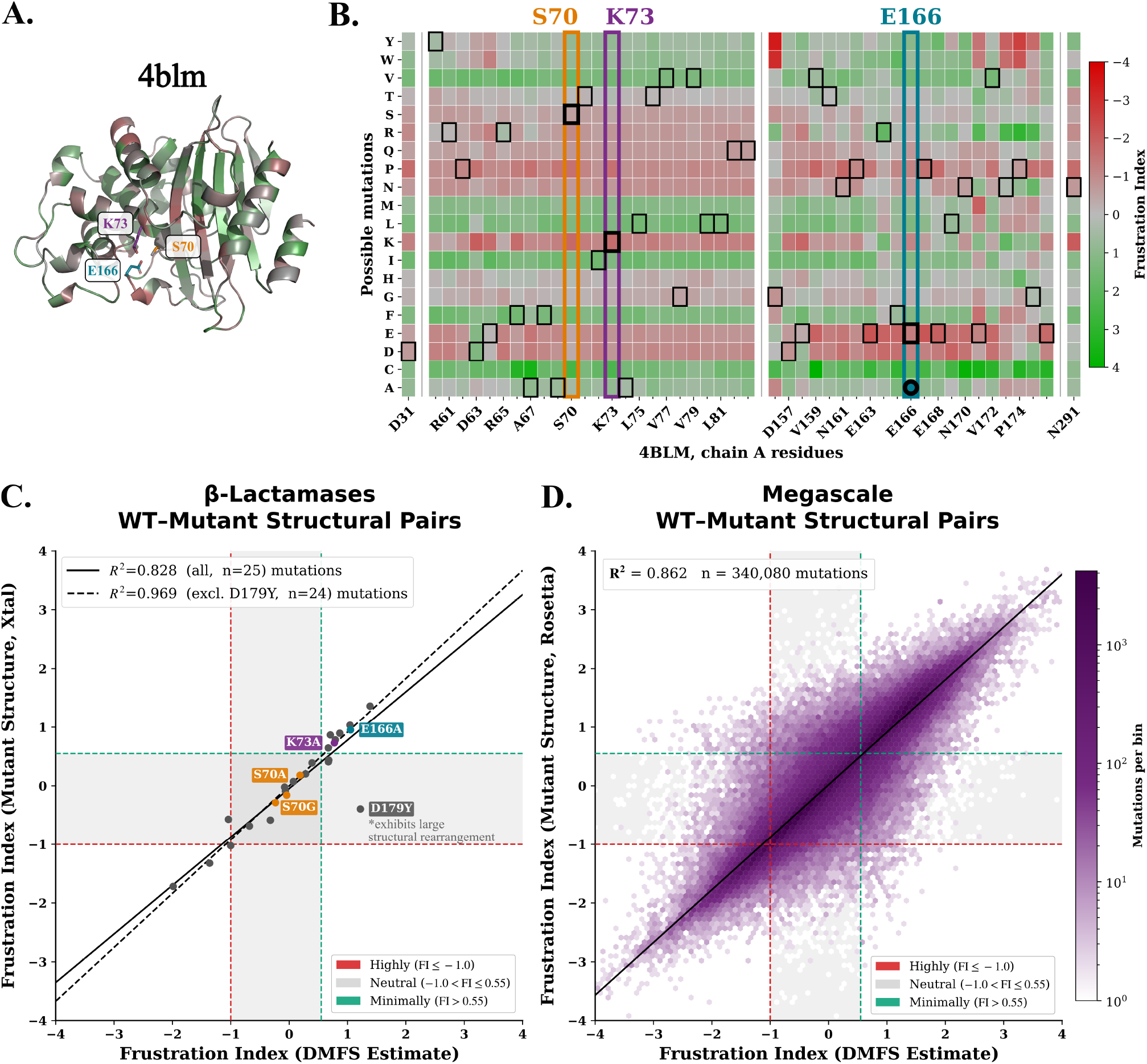
Deep mutational frustration scanning of local energetic frustration. PyTorch Frustratometer enables rapid deep mutational frustration scanning (DMFS) from a single wild-type structure, estimating the local frustration associated with all possible amino acid substitutions. (A) Single-residue frustration mapped onto *Bacillus licheniformis β*-lactamase (PDB ID: 4BLM, chain A), highlighting the catalytic residues S70, K73 and E166. (B) DMFS frustration matrix for 4BLM, chain A, with truncated residue windows (“…”) to highlight known mutations. Black boxes indicate the native amino acid at each position, while the circle indicates an experimentally characterized mutation in *B. licheniformis* at E166A (Knox et al., 1993). Wild-type catalytic residues are highly frustrated (↓ frustration index, ↑ red); hydrophobic substitutions generally shift these positions toward minimal frustration (↑ frustration index, ↑ green), whereas several polar, charged and small/flexible substitutions retain frustration levels closer to the wild type (↓ frustration index, ↑ red). (C) Comparison between frustration indices estimated by DMFS from the wild-type *β*-lactamase structures and those calculated from paired experimentally determined *β*-lactamase mutant structures from *K. pneumoniae, B. licheniformis, E. coli, S. aureus*, and *E. cloacae* (n = 25; R^2^ = 0.828), where highlighted mutations are shared by panels A, B, C. We show mutations in catalytic residues: S70 in yellow (S70G and S70A (Stojanoski et al., 2016) from *K. pneumoniae*); K73 in teal (K73A (Hinchliffe et al., 2022) from *K. pneumoniae)* and E166 in teal (E166A from *B. licheniformis* (Knox et al., 1993)) as well as mutations from non-catalytic residues from the other species in gray. One mutant exhibiting substantial structural rearrangements (D179Y) is shown to highlight the limits of this algorithm, which when removed, raises the correlation (n = 24; R^2^ = 0.969) (Supp. Table 1). (D) Large-scale validation using the MegaScale dataset, comparing DMFS estimates from 298 wild-type structures with frustration indices calculated from 340,080 corresponding Rosetta-modelled single-mutant structures (R^2^ = 0.862).

To further test the ability of DMFS to predict mutation-induced changes in frustration, we compared DMFS estimates with frustration values calculated from a dataset of publicly available Rosetta mutant models comprising 340,080 single-amino-acid point mutation models across 298 proteins (Dieckhaus and Kuhlman, 2025). This allowed us to systematically compare DMFS values calculated directly from wild-type structures with those obtained from the corresponding Rosetta-modeled mutant structures. The two estimates showed strong agreement (R^2^ = 0.862; Fig. 2D), further supporting the new DMFS module as a scalable approximation of how local energetic frustration changes upon single amino acid substitutions.

### Massive performance gains

We next evaluated the computational performance of PyTorch Frustratometer using proteins spanning a broad range of sizes and compared runtimes with the original FrustratometeR implementation. Across all benchmarks, PyTorch Frustratometer substantially reduced execution times while producing consistent values for the different frustration indices, within expected errors (Fig. 3). The largest performance gains were observed for large proteins and batch calculations, where vectorized tensor operations and GPU acceleration efficiently parallelize the evaluation of native and decoy interactions. For the three conventional frustration modes (Fig. 3A-C), PyTorch Frustratometer achieved wall clock speedups of °70× (configurational) to over 13,000*×* (mutational) depending on protein size and frustration mode, while substantially larger gains were observed for DMFS, reaching °137,000× acceleration (Fig. 3D). GPU acceleration also enabled the analysis of very large macromolecular assemblies that could not be processed with FrustratometeR because of memory limitations or due to the required legacy PDB format input, as indicated by the largest successfully processed structures by all three modes and DMFS in Fig. 3A–D, that had no memory or format issues. At this limit, for example, analysis of the Squid Hemocyanin structure (PDB ID: 4YD9 chain A, 1,656 residues) required 24.72, 322.80, and 5.27 seconds (wall clock) for single-residue, mutational, and configurational frustration, respectively, compared with only 0.052, 0.062, and 0.049 seconds using GPU-accelerated PyTorch Frustratometer (Supp. Table 2). For the same protein, the difference was even more pronounced for DMFS, which required approximately 3 hours (10,680 seconds) with FrustratometeR but only 78 milliseconds with GPU-accelerated PyTorch Frustratometer, corresponding to a speedup of °137,000×. The most extreme examples to showcase the enhanced scalability offered by PyTorch Frustratometer include the 13,364-residue LP ring (PDB ID: 8WHT), analyzed for single-residue frustration in just 0.976 seconds, and the 45,108-residue flagellar motor-hook complex (PDB ID: 7CGO), analyzed for mutational frustration in 4.951 seconds (Fig. 3E,F). Note that all timings are wall clock time, reported as the median of 3 repetitions, excluding GPU warm up times. These performance gains make routine frustration analysis of proteome-scale structural collections, extensive molecular dynamics trajectories, and large protein design campaigns computationally feasible.

**Figure 3.**
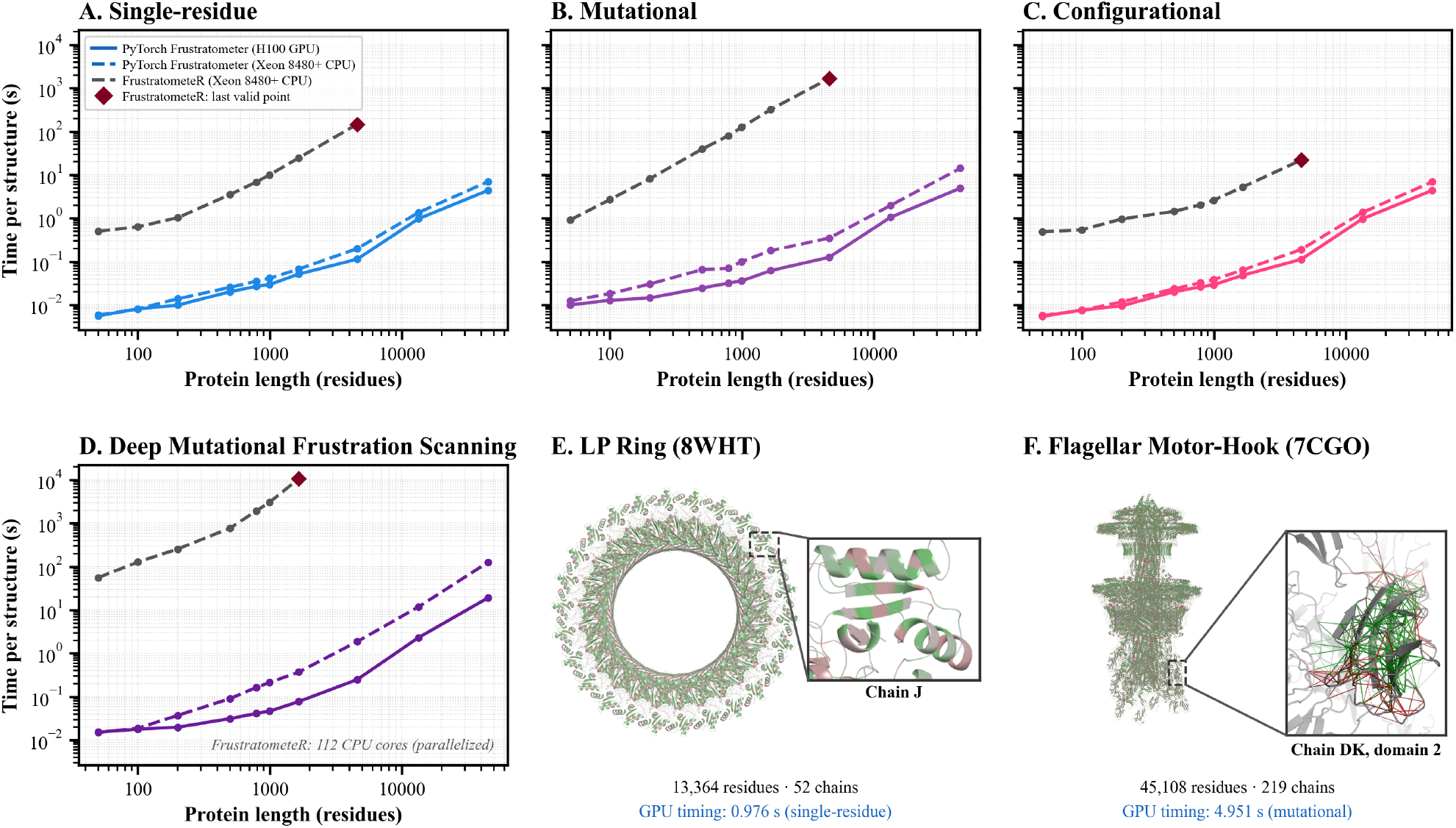
Computational performance and scalability of PyTorch Frustratometer. Execution time per structure as a function of protein length for **(A)** single-residue, **(B)** mutational, **(C)** configurational, and **(D)** deep mutational frustration scanning (DMFS) frustration analyses. PyTorch Frustratometer performance using an NVIDIA H100 GPU (solid lines) and an Intel Xeon 8480+ CPU (dashed colored lines) is compared with FrustratometeR running on an Intel Xeon 8480+ CPU (grey dashed lines) using minimum sequence separation of 3, electrostatics enabled, and for configurational mode, 2000 random decoys. Red diamonds indicate the largest structures successfully processed by FrustratometeR with no out of memory or format parsing issues. For DMFS, FrustratometeR calculations were parallelized across 112 CPU cores, using a custom parallelizable version. **(E**,**F)** Examples illustrating the scalability of GPU-accelerated calculations to very large macromolecular assemblies that FrustratometeR could not handle: **(E)** the LP ring (PDB ID: 8WHT; 13,364 residues, 52 chains), analyzed for single-residue frustration in 0.976 s (GPU) and 1.359 s (CPU); and **(F)** the flagellar motor-hook complex (PDB ID: 7CGO; 45,108 residues, 219 chains), analyzed for mutational frustration in 4.951 s (GPU) or 14.22 s (CPU).

## Discussion

PyTorch Frustratometer substantially expands the scale at which local energetic frustration can be analyzed while preserving the conceptual framework and interpretation of previous FrustratometeR implementations. Its near-perfect numerical agreement with FrustratometeR, despite the random sampling involved in the calculation of the decoy distributions, shows that the computational gains from tensorization and GPU acceleration do not come at the expense of methodological consistency.

This new implementation enables analyses that were previously computationally prohibitive. In particular, deep mutational frustration scanning provides a rapid approximation of how substitutions reshape the local energetic environment without requiring explicit structural models for every mutant. The agreement with experimentally determined mutant structures supports this interpretation and opens opportunities for integrating frustration analysis with large-scale mutational and protein-design studies.

The fast expansion of experimentally determined and predicted structures is shifting structural bioinformatics from the analysis of individual proteins toward datasets containing thousands to millions of models. By reducing calculations that previously required hours to seconds or minutes and supporting parallel CPU/GPU execution, PyTorch Frustratometer makes local frustration analysis compatible with proteome-scale datasets, large conformational ensembles, and high-throughput protein design pipelines. This increased scalability is particularly timely as local energetic frustration is increasingly being used to connect protein evolution, conformational dynamics, and design, while also being integrated with modern AI-based approaches (Simonetti et al., 2026).

The recent widespread adoption of AI-based tools, such as FrustrAI-seq, enables rapid frustration predictions at unprecedented scales (Leusch et al., 2026); however, physics-based approaches such as PyTorch Frustratometer remain essential for providing interpretable, mechanistic insight into the energetic interactions underlying these predictions, complementing the capabilities of AI-based methods. In combination, these developments provide a foundation for studying energetic frustration at the scale of the known protein structure universe.

### Methods

Most of the code is a direct port of the AWSEM (Davtyan et al., 2012) C++ code into PyTorch. We have re-implemented the water, burial, and electrostatic energy terms exactly as they appear in the original AWSEM, using all of the same parameters. Because PyTorch allows for batched operations, we were able to score all of the generated decoys together in a single computation, resulting in faster speeds. FrustratometeR (Jenik et al., 2012; Parra et al., 2016; Rausch et al., 2021) is an R wrapper for AWSEM handling IO, parameter selection, and running the C++ AWSEM binary. We now use PyTorch Lightning to handle IO, parameter selection, and inference, borrowing the protein data structures and atom placement geometry from AlphaFold (Jumper et al., 2021), OpenFold (Ahdritz et al., 2023), and Frame2seq (Akpinaroglu et al., 2023). Please see the Supplementary Methods for a detailed description of how memory is managed for GPU acceleration as well as how input protein structures are processed.

For each mode we calculate the frustration as:

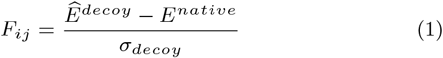

representing a z-score where 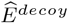 is the mean of the decoy energy distribution, *E*^*native*^ is the native energy, and *σ*_*decoy*_ is the standard deviation of the decoy energy distribution. The decoy distribution is estimated either by doing an exhaustive enumeration for single-residue mode and mutational mode, *or* by creating 100,000 random decoy structures for each residue pair in configurational mode. Frustration indices typically fall within approximately *±*4 standard deviations, with values beyond this range being rare in natural proteins (Ferreiro et al., 2007, 2014; Parra et al., 2016). We describe how frustration is calculated for each mode in the Supplementary Methods.

### Pairwise Smooth Frustration

We also implement the smooth frustration index introduced by Guan et al. (2024), which aids in identifying highly frustrated domain interface contacts in a clearer way than the mutational mode. Instead of sampling decoys, the smooth index is derived directly from the single-residue frustration indices:

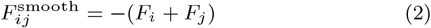

where *F*_*i*_ and *F*_*j*_ are the native single-residue frustration indices at positions *i* and *j*. Classification uses the smooth-frustration thresholds, which are inverted from the original mutation mode threshold, introduced by Guan et al. (2024) (minimally frustrated: F *< −*1.2; highly frustrated: F *>* 0.8). (Supp. Fig. 3)

### Frustration Classification

Frustration indices are classified into three categories using mode-specific thresholds. For single-residue frustration, residues are classified as highly frustrated when FI *≤ −*1.0, neutrally frustrated when *−*1.0 *<* FI *≤* 0.55, and minimally frustrated when FI *>* 0.55. For mutational and configurational frustration, interactions are classified as highly frustrated when FI *≤ −*1.0, neutrally frustrated when *−*1.0 *<* FI *≤* 0.78, and minimally frustrated when FI *>* 0.78. These thresholds follow the original FrustratometeR convention (Ferreiro et al., 2007; Parra et al., 2016). For smooth frustration, the direction of the index is inverted: interactions are classified as highly frustrated when FI *>* 0.8, neutrally frustrated when *−*1.2 *<* FI *≤* 0.8, and minimally frustrated when FI *≤ −*1.2, following the thresholds defined by Guan et al. (2024).

### Mutational Scanning

In FrustratometeR, mutation scanning requires O(L *×* 20) independent AWSEM runs per protein, one for each amino acid substitution at each position, with missing atoms for glycine mutations filled in with MODELLER (Webb and Sali, 2016).

We instead evaluate all L *×* 20 amino acid substitutions in a single batched operation using the ΔE approach that we use to compute the decoy energies for single-residue frustration (see Supplementary Methods for Single Residue Frustration; equation S2), producing a complete [L *×* 20] frustration index table reducing lookup times to O(1).

Glycine requires two corrections since it uses C*α* instead of C*β* as its centroid:

- G to X: for native glycine positions, we place a virtual C*β* using CHARMM36 (Huang and MacKerell, 2013) internal coordinates and compute absolute energies from scratch in vC*β* geometry, since switching centroids affects the full decoy distribution.
- X to G: for glycine substitutions at non-glycine positions, we compute a separate all-C*α* decoy distribution and extract the GLY frustration index from it.

All other positions use the standard ΔE approach.

### Datasets

We benchmarked PyTorch Frustratometer against the CATH v4.4 S40 dataset, a nonredundant dataset of 34,653 CATH domain structures, constructed from representative structures after clustering CATH at 40% sequence identity (Waman et al., 2025). We further process each domain structure using pdb tools (Rodrigues et al., 2018) by (i) selecting the highest occupancy altlocs (note this is different than the PyTorch Frustratometer default option as this is just an internal validation set); (ii) fixing insertion codes by increasing the residue number and shifting downstream residue numbers; (iii) renumbering all residues from 1; and finally

(iv) renumbering all atoms from 1. Finally, we ran the entire dataset through FrustratometeR for single-residue, mutational, and configurational modes using default parameters: SeqDist=3, and Electrostatics K=4.15. Finally, all PyTorch Frustratometer runs were performed using the same default parameters as FrustratometeR (min-seq-sep=3, burial-min-sep=3 [to match FrustratometeR, but note the discrepancy in the Default Parameters section], and electrostatics *on* with k=4.15) along with the completed PDB files from FrustratometeR to ensure they were parsing the same PDB coordinates.

### Default values for user tunable parameters

After benchmarking different implementations, we have decided on a set of recommended parameter values for all frustration calculations going forward that we make as default:

**--min-seq-sep:** Minimum sequence separation |i-j| *>*= value for a contact to be considered. **Default value: 3**. In FrustratometeR, this parameter was called “SeqDist.” Previous implementations used 12, which is not recommended anymore. Note that this does not affect the distance used for burial which is set to --burial-min-sep *>* 1.

**--burial-min-sep:** Minimum sequence separation |i-j| *>* value for residue density (rho) calculation. **Default value: 1**, i.e. there needs to be at least 1 residue between residues i and j. In some recent versions of FrustratometeR, this value was set to 3 or 12 to match “--min-seq-sep”. Our comparisons between DMFS and values explicitly calculated on mutant structural models showed that the best value is 1 as it leads to the highest correlation. This value should not be changed and is hidden from the list of arguments.

**--decoys:** Number of decoys for configurational frustration index calculation. **Default: 100 000**. Fewer decoys would make the tool faster but would lead to lower convergence across independent runs on the same data. This parameter can be decreased to as few as 2,000 decoys without substantially compromising accuracy.

**--cutoff:** Contact distance cutoff in Å. **Default: 9.5**. Standard across all FrustratometeR and AWSEM implementations. We do not recommend changing this value.

**--electrostatics:** Controls electrostatic terms. **Default: True**, that sets the electrostatics constant to its default value k=4.15 which corresponds to an aqueous environment. Other values for k can be used by setting --electrostatics directly to the k constant value (e.g. -- electrostatics 7). Set --electrostatics false to turn off the electrostatics term. All previous versions run without electrostatics by default.

## Acknowledgements

Authors thank MareNostrum5 for computational resources at the BSC. EJD is funded by the DeCYPher Horizon Europe Project (grant number 10108178). RGP is funded by the Ramon y Cajal program (RYC2023-043825-I) and the MEGAFrustratEDS grant (PID2024-159128OA-I00). FLS is a fellow within the “Generación D” initiative, Red.es, Ministerio para la Transformación Digital y de la Función Pública, for talent attraction (C005/24-ED CV1). DUF is supported by the Consejo de Investigaciones Científicas y Técnicas (CONICET); CONICET Grant PIP2022-2024—11220210100704CO and Universidad de Buenos Aires grant UBACyT 20020220200106BA. Work at Rice University was supported by NSF grant PHY-2019745 for the Center for Theoretical Biological Physics, NSF grant CHE-2136142 fellowship from the Molecular Software Institute and the Bullard - Welch Chair at Rice University, Grants-0016.

