## Supplementary Material for "PyTorch Frustratometer: An Ultra-fast, Parallelized Tool for Proteome-Scale Local Energetic Frustration Analysis"

### Supplementary Figures and Tables

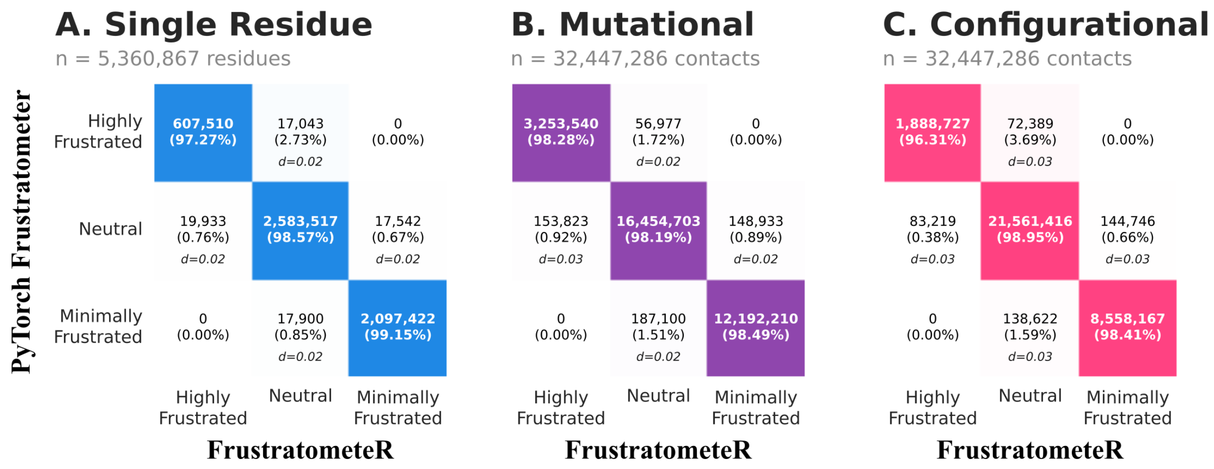


**Supp. Fig. 1. Confusion matrices of comparing PyTorch Frustratometer classification to FrustratometeR.** For each mode (A-C), PyTorch Frustratometer predicts the correct class (Highly, Neutrally, or Minimally Frustrated) $> 96\%$ of the time, where mismatches occur near the threshold boundaries, compared to the original FrustratometeR on the CATH v4.4 S40 dataset. We classify raw frustration indices using the given thresholds: single-residue (Highly: $FI \leq -1$; Neutral: $-1 < FI \leq 0.55$, Minimally: $FI > 0.55$), mutational (Highly: $FI \leq -1$; Neutral: $-1 < FI \leq 0.78$, Minimally: $FI > 0.78$), configurational (Highly: $FI \leq -1$; Neutral: $-1 < FI \leq 0.78$, Minimally: $FI > 0.78$). For mismatches off of the diagonal, we show the average distance (d) to the known threshold boundary for that wrongly predicted class. All mismatches are less than 0.03 FI units to the boundary.

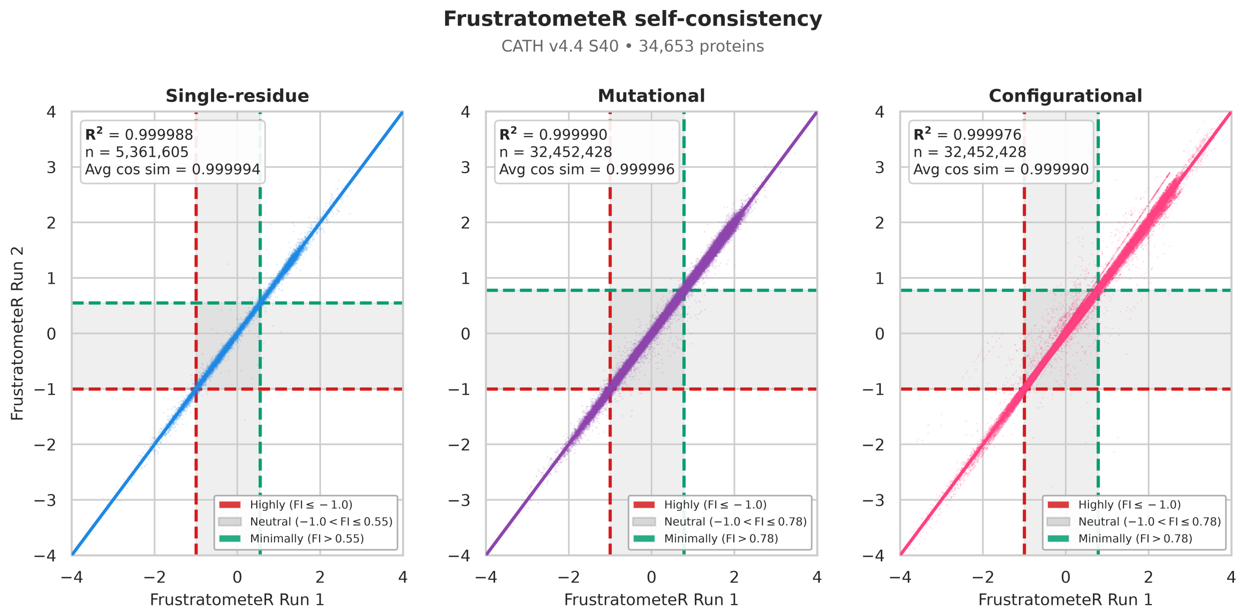


**Supp. Fig. 2. Self-consistency of FrustratometeR across repeated runs.** Using the CATH v4.4 S40 dataset, we note that the original FrustratometeR has some discrepancies between multiple runs, due to minor random sampling effects. This shows that minor variations between PyTorch Frustratometer and the original FrustratometeR are within the expected error range.

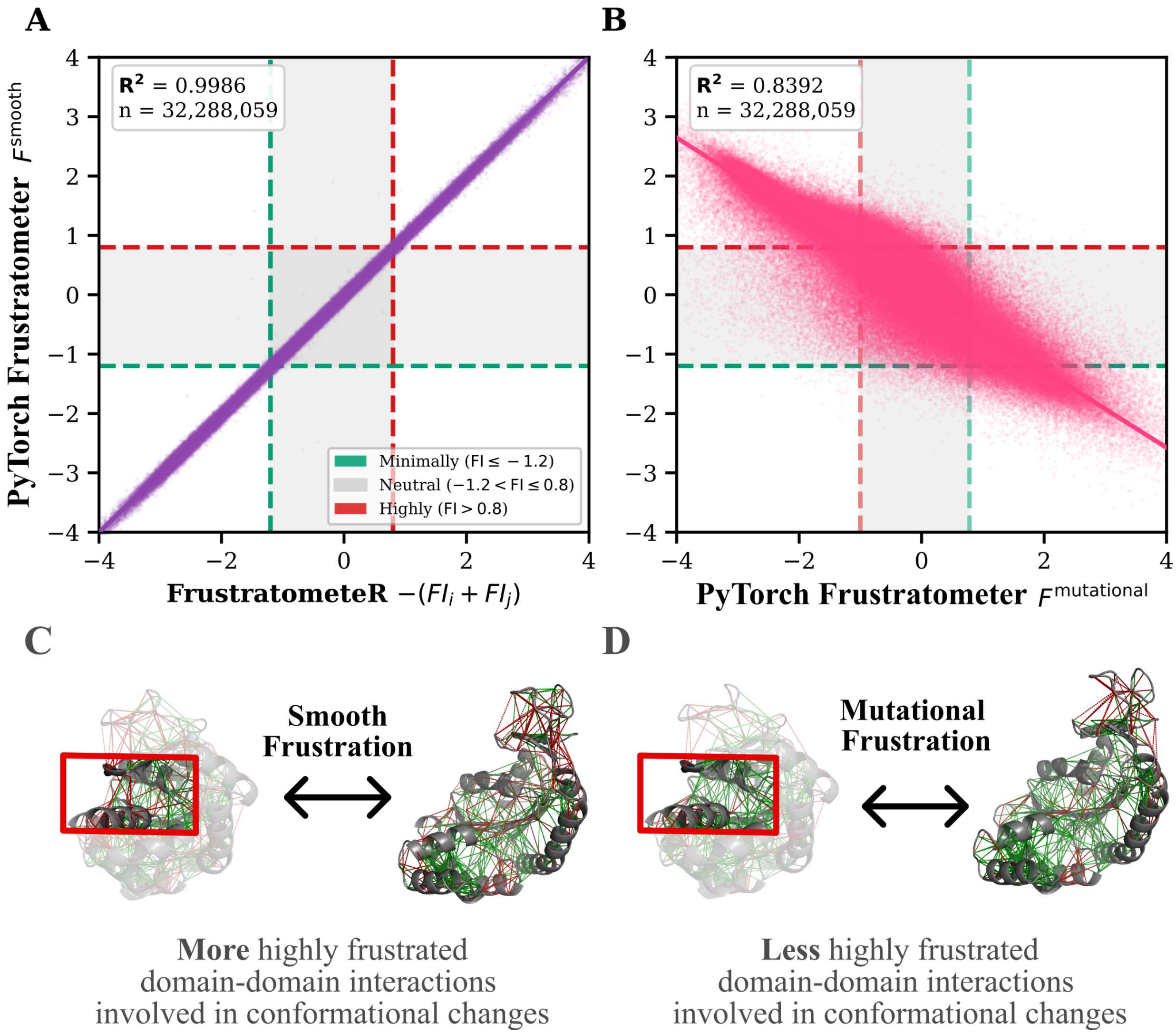


**Supp. Fig. 3. Smooth Frustration is a new frustration mode.** (A) The new mode performs $F^{smooth}$ = - (Fi + $F_j$) from single-residue frustration indices. When performing the same calculation with FrustratometeR single-residue frustration indices, there is a strong agreement ($R^2$=0.9994) compared to the CATH v4.4 S40 dataset. (B) The smooth mode shows a strong inverse relationship with the mutational mode ($R^2$=0.8383), showing good agreement but captures different information. In both panels, we used min-seq-sep=3, decoys=1000, electrostatics_k=4.15, and burial-min-sep=3 to match the original FrustratometeR. (C) Smooth frustration identifies more highly frustrated contacts in domain-domain interactions involved in conformational changes, than (D) mutational frustration, both represented by the adenylate kinase (ADK) structures in closed conformation (PDB ID: 1AKE chain A) and open conformation (PDB ID: 4AKE chain A).

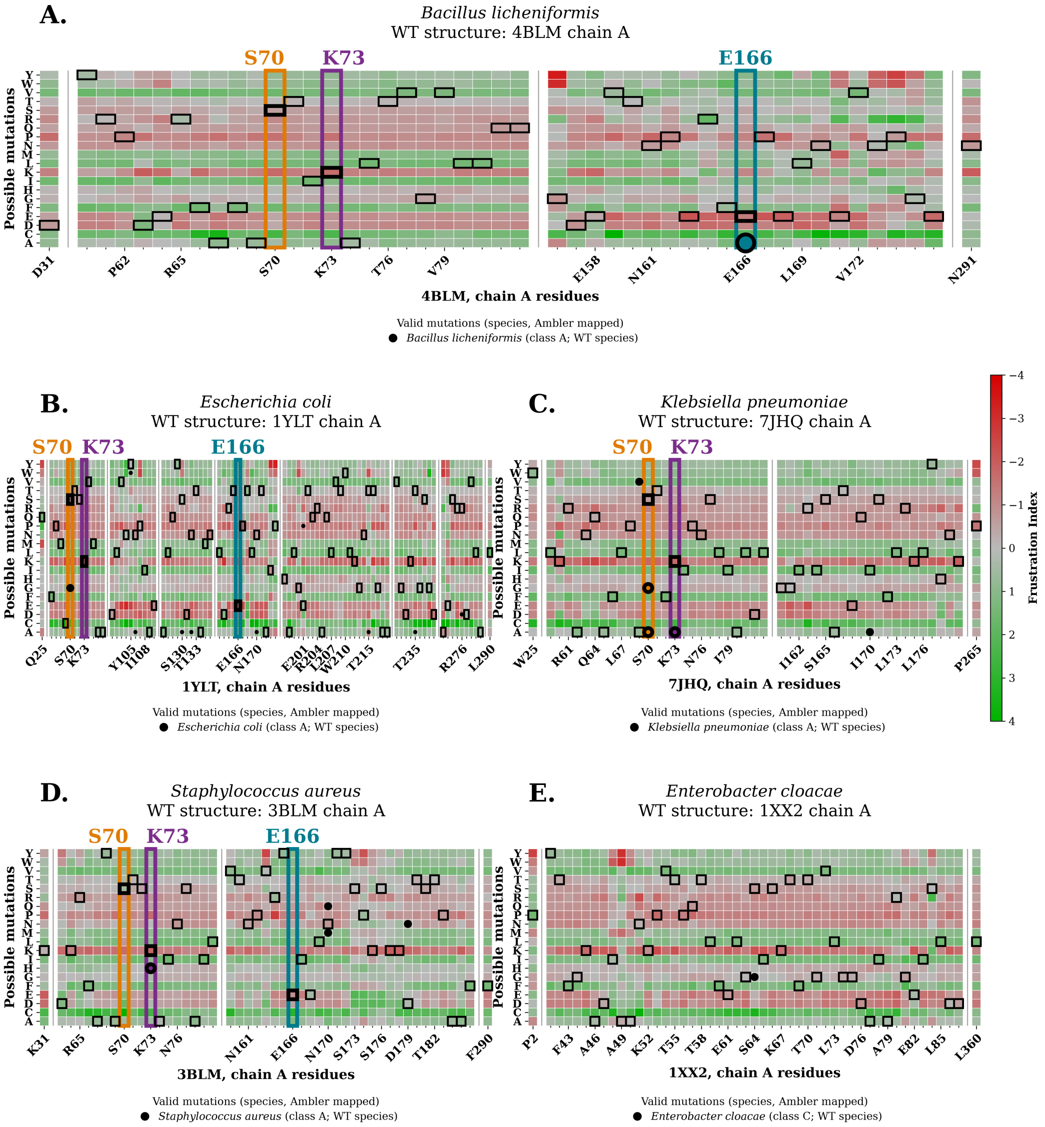


**Supp. Fig. 4. DMFS frustration matrices for $\beta$-lactamase from different species.** We show known mutations with pairs of experimentally calculated wild type and mutant structures as circles, which also appear in Fig. 2B and 2C. Black boxes represent the wild type amino acid for each representative structure. (A) DMFS frustration matrix for a Class A $\beta$-lactamase from *B. licheniformis* (PDB ID: 4BLM, chain A), with 1 circle marking the E166A mutation that has WT-Mutant structural pair. (B) DMFS frustration matrix for a Class A $\beta$-lactamase from *E. coli* (PDB ID: 1YLT, chain A), with 11 circles marking mutations that have WT-Mutant structural pairs from itself and *other* CTX-M-14 $\beta$-lactamases (same class as 1YLT) from *E. coli*, as well as mutations from TEM-1 $\beta$-lactamases from *E. coli*, mapped using Ambler notation. (C) DMFS frustration matrix for a Class A $\beta$-lactamase from *K. pneumonia* (PDB ID: 2OV5, Chain A), with 4 circles marking mutations that have WT-Mutant structural pairs from itself and *other SHV* $\beta$-lactamases from *K. pneumonia,* excluding two pairs that come from Class D $\beta$-lactamases. (D) DMFS frustration matrix for a Class A $\beta$-lactamase from a *S. aureus* (PDB ID: 3BLM, Chain A), with 4 circles marking mutations that have WT-Mutant structural pairs from itself and *other* Class A $\beta$-lactamases from *S. aureus*. (E) DMFS frustration matrix for a Class C $\beta$-lactamase from a *E. cloacae* (PDB ID: 1XX2, Chain A), with 1 circle marking a mutation that has a WT-Mutant structural pair. A full list of mutations included in these DMFS frustration matrices (A–E) and used in Fig. 2C is provided in Supp. Table 1.

| Mutation | WT PDB | Mutant PDB | WT Frustration | Mutant Frustration | DMFS-estimated Mutant Frustration | WT Organism | Mutant Organism |
|---|---|---|---|---|---|---|---|
| **S70A** | 7JHQ | 5HAP | -0.599 | 0.179 | 0.186 | *K. pneumoniae* | *K. pneumoniae* |
| **K73A** | 4HBT | 7QQC | -1.543 | 0.73 | 0.776 | *E. coli* | *K. pneumoniae* |
| **E166A** | 1BTL | 1MBL | -1.51 | 0.952 | 1.051 | *E. coli* | *B. licheniformis* |
| **S235A** | 1XPB | 1ESU | -0.61 | 0.437 | 0.678 | *E. coli* | *E. coli* |
| **N170M** | 3BLM | 1KGE | -0.775 | 0.782 | 0.792 | *S. aureus* | *S. aureus* |
| **N170Q** | 3BLM | 1KGF | -0.775 | -0.685 | -0.681 | *S. aureus* | *S. aureus* |
| **M69V** | 1SHV | 3V5M | 0.851 | 1.361 | 1.386 | *K. pneumoniae* | *K. pneumoniae* |
| **L201P** | 1XPB | 3CMZ | 1.133 | -1.719 | -1.987 | *E. coli* | *E. coli* |
| **M182T** | 1BTL | 1JWP | 0.78 | -0.023 | -0.078 | *E. coli* | *E. coli* |
| **N276D** | 1BTL | 1CK3 | -1.045 | -1.323 | -1.362 | *E. coli* | *E. coli* |
| **K73H** | 3BLM | 1DJA | -1.778 | 0.39 | 0.392 | *S. aureus* | *S. aureus* |
| **N106A** | 1YLT | 8DOE | -0.851 | 0.866 | 0.706 | *E. coli* | *E. coli* |
| **S130A** | 1YLT | 8DOD | 0.143 | 0.071 | 0.072 | *E. coli* | *E. coli* |
| **N132A** | 1YLT | 8ELB | -0.816 | 0.644 | 0.671 | *E. coli* | *E. coli* |
| **T215A** | 1YLT | 8DON | -0.264 | -0.572 | -1.041 | *E. coli* | *E. coli* |
| **T235A** | 1YLT | 8DP4 | 0.362 | 0.202 | 0.282 | *E. coli* | *E. coli* |
| **Y105W** | 1YLT | 7UON | 0.082 | 0.408 | 0.673 | *E. coli* | *E. coli* |
| **N170A** | 1YLT | 8DPQ | -0.611 | 1.033 | 1.043 | *E. coli* | *E. coli* |
| **N170A** | 5UL8 | 6XD5 | -1.028 | 0.895 | 0.867 | *K. pneumoniae* | *K. pneumoniae* |
| **D179N** | 3BLM | 1BLP | 0.254 | -0.586 | -0.322 | *S. aureus* | *S. aureus* |
| **D179N** | 2OV5 | 7TB7 | -1.394 | -1.024 | -1.001 | *K. pneumoniae* | *K. pneumoniae* |
| **D179Y** | 2OV5 | 7TBX | -1.394 | -0.395 | 1.221 | *K. pneumoniae* | *K. pneumoniae* |
| **S64G** | 1XX2 | 5HAI | -0.693 | -0.085 | -0.065 | *E. cloacae* | *E. cloacae* |
| **S70G** | 1ZG4 | 1ZG6 | -0.365 | -0.16 | -0.042 | *E. coli* | *E. coli* |
| **S70G** | 7JHQ | 5HAQ | -0.599 | -0.287 | -0.234 | *K. pneumoniae* | *K. pneumoniae* |

**Supp. Table 1. $\beta$-lactamase mutations used for validation of deep mutational frustration scanning (DMFS) frustration estimates.** For each mutation, the table reports the experimentally determined wild-type (WT) and mutant structures (PDB IDs), the single-residue frustration index calculated directly from the WT structure (**WT Frustration**), the frustration index calculated directly from the experimentally determined mutant structure (**Mutant Frustration**), and the frustration index predicted for the mutation from the WT structure using DMFS (**DMFS-estimated Mutant Frustration**). WT and mutant organisms are also indicated. All mutations were included in the main correlation analysis (Fig. 2C), whereas D179Y is shown because it highlights a core limitation of the algorithm in which the structure undergoes substantial backbone rearrangements upon mutation, which is visible in its experimentally determined mutant structure. The table therefore provides the individual structural pairs and frustration values underlying the validation shown in Fig. 2C.

| PDB | 1A92 C | 1A30 A | 1A0F A | 1A8H A | 1OH6 A | 3W5B A | 4YD9 A | 9FQR HL | 8WHT | 7CGO |
|---|---|---|---|---|---|---|---|---|---|---|
| **n (residues)** | **50** | **100** | **201** | **500** | **788** | **997** | **1656** | **4588** | **13364** | **45108** |
| **SR GPU (s)** | 0.005604 | 0.008116 | 0.009979 | 0.02031 | 0.02688 | 0.02969 | 0.05179 | 0.1143 | 0.9761 | 4.42 |
| **SR CPU (s)** | 0.005863 | 0.008136 | 0.01394 | 0.02564 | 0.03532 | 0.0414 | 0.06749 | 0.1998 | 1.359 | 7.019 |
| **SR FR (s)** | 0.506 | 0.633 | 1.035 | 3.543 | 6.835 | 10.09 | 24.72 | 147.7 | -- | -- |
| **Mut GPU (s)** | 0.0101 | 0.01284 | 0.01462 | 0.02469 | 0.03201 | 0.03593 | 0.06235 | 0.1265 | 1.055 | 4.951 |
| **Mut CPU (s)** | 0.01251 | 0.01835 | 0.03051 | 0.06588 | 0.07045 | 0.1005 | 0.1801 | 0.3509 | 1.982 | 14.22 |
| **Mut FR (s)** | 0.914 | 2.703 | 8.185 | 39.45 | 80.11 | 126 | 322.8 | 1683 | -- | -- |
| **Conf GPU (s)** | 0.005465 | 0.007643 | 0.009618 | 0.02031 | 0.02609 | 0.02931 | 0.04858 | 0.1134 | 0.9745 | 4.418 |
| **Conf CPU (s)** | 0.005709 | 0.007586 | 0.01188 | 0.02374 | 0.0325 | 0.03865 | 0.0643 | 0.1913 | 1.367 | 6.931 |
| **Conf FR (s)** | 0.489 | 0.544 | 0.968 | 1.447 | 2.048 | 2.619 | 5.271 | 22.09 | -- | -- |
| **DMFS GPU (s)** | 0.01535 | 0.01799 | 0.01999 | 0.03152 | 0.04145 | 0.04666 | 0.07784 | 0.2502 | 2.322 | 19.28 |
| **DMFS CPU (s)** | 0.01505 | 0.01893 | 0.03743 | 0.0911 | 0.165 | 0.214 | 0.373 | 1.877 | 11.88 | 127.4 |
| **DMFS FR (s)** | 56.09 | 128 | 257.5 | 778.1 | 1930 | 3081 | $1.07\times10^4$ | --* | -- | -- |

**Supp. Table 2. Raw timing benchmarks across protein sizes, frustration modes, and hardware.** For each protein structure (PDB), the number of residues (**n**) and wall-clock execution time in seconds (**s**) are reported for single-residue (**SR**), mutational (**Mut**), configurational (**Conf**), and deep mutational frustration scanning (**DMFS**) frustration calculations. **GPU** and **CPU** indicate PyTorch Frustratometer calculations performed using GPU and CPU hardware, respectively, whereas **FR** indicates calculations performed with the original FrustratometeR implementation. PyTorch Frustratometer achieves approximately 70–13,000× acceleration on GPU for conventional frustration modes, depending on protein size and calculation mode, with substantially larger gains for DMFS. —* indicates that FrustratometeR crashed due to out-of-memory (OOM) errors, whereas — indicates that FrustratometeR could not parse the input structure because of limitations of the legacy PDB format for very large systems. This table provides the raw timing values underlying the performance and scalability comparisons shown in Fig. 3.

### Supplementary Methods

#### Calculation of frustration in each mode

The single-residue, mutational and configurational frustration indices are calculated as follows.

##### Single-Residue

We compute the energy for all 20 amino acid type substitutions at each position exactly, weighted by sequence composition, instead of sampling random decoys. This is equivalent to the limit of sampling infinite decoys (lim D→∞) of the C++ random sampling approach, which is both faster and more stable. We first calculate the native energy at each position:

$$\begin{aligned} E^{\text{native}}(i) = E^{\text{burial}}(i) \\ + \sum_{\substack{k \neq i \\ r_{ik} < r_c}} \left[ E^{\text{water}}(i,k) + E^{\text{elec}}(i,k) \right] \end{aligned} \tag{S1}$$

where $i$ is an amino acid position and $k$ is a residue that has a distance to i ($r_{ik}$) that is within the contact cutoff ($r_c$ *; we use 9.5 Å*). If the electrostatic term is disabled, $E^{\text{elec}}$ will be zero.

For each position $i$ with native amino acid type $a$, we compute the decoy energy for substitution $a'$ as a delta from the native:

$$\begin{aligned} E^{decoy}(i,a') = E^{native}(i) + \Delta E^{burial}(i,a') \\ + \Delta E^{water}(i,a') + \Delta E^{elec}(i,a') \end{aligned} \tag{S2}$$

where ΔE terms are differences between the decoy and native contributions at position i, with all neighbour positions and densities held fixed. The decoy mean and standard deviation are weighted by sequence composition f(a′), the fraction of amino acid type a′ in the sequence:

$$\widehat{E}^{decoy,sequence}(i) = \sum_{a'} f(a')\, E^{decoy}(i,a') \tag{S3}$$

$$\sigma^{decoy,sequence}(i) = \sqrt{\sum_{a'} f(a') \left( E^{decoy}(i,a') - \widehat{E}^{decoy,sequence}(i) \right)^2} \tag{S4}$$

If a residue type is missing from the original sequence, it will not be sampled from. However, you can change the distribution type to allow pseudo-counts, which will sample missing residue types.

##### Mutational

For every native contact pair ($i$, $j$) the interaction energy includes direct terms plus contributions from all background residues $k$ within 9.5 Å of either $i$ or $j$:

$$
\begin{aligned}
E^{\text{native}}(i,j) = {} & E^{\text{water}}(i,j) + E^{\text{elec}}(i,j) + E^{\text{burial}}(i) + E^{\text{burial}}(j) \\
& + \sum_{\substack{k\neq i,j \\ r_{ik}<r_c}} \Big[E^{\text{water}}(i,k) + E^{\text{elec}}(i,k)\Big] \\
& + \sum_{\substack{k\neq i,j \\ r_{jk}<r_c}} \Big[E^{\text{water}}(j,k) + E^{\text{elec}}(j,k)\Big]
\end{aligned}
\tag{S5}
$$

where $i$ and $j$ are the residues involved in a contact (d<$r_c$=9.5) and $k$ is a residue that has a distance to $i$ ($r_{ik}$) or j ($r_{jk}$) that is within the contact cutoff ($r_c$ *; we use 9.5 Å*). If the electrostatic term is disabled, $E^{\text{elec}}$ will be zero.

We precompute, once per structure, the burial energy, background water energy, and if enabled, electrostatic field, for all 20 amino acid types at every position and edge endpoint. This lets us score each decoy as a direct table lookup (O(1)), allowing us to evaluate all 400 pairs per contact without recomputing any background sums.

We then compute the decoy energies for all 400 amino acid pair substitutions (*a', b'*) at each contact exactly, weighted by sequence composition, instead of sampling random decoys. This is equivalent to the limit of sampling infinite decoys (lim D→∞) of the C++ random sampling approach, which removes sampling noise. In this mode, backbone positions and densities are fixed. Each decoy energy is:

$$
\begin{aligned}
E^{\text{decoy}}(i,a',j,b') = {} & E^{\text{water}}(i,a',j,b') + E^{\text{elec}}(i,a',j,b') \\
& + E^{\text{burial}}(i,a') + E^{\text{burial}}(j,b') \\
& + \sum_{\substack{k\neq i,j \\ r_{ik}<r_c}} \Big[E^{\text{water}}(i,a',k) + E^{\text{elec}}(i,a',k)\Big] \\
& + \sum_{\substack{k\neq i,j \\ r_{jk}<r_c}} \Big[E^{\text{water}}(j,b',k) + E^{\text{elec}}(j,b',k)\Big]
\end{aligned}
\tag{S6}
$$

where *a'* and *b'* are the decoy residue types at positions $i$ and $j$, and $k$ is a residue that has a distance to $i$ ($r_{ik}$) or j ($r_{jk}$) that is within the contact cutoff ($r_c$ *; we use 9.5 Å*). The decoy mean and standard deviation are weighted by the probability $f(a')f(b')$ of drawing the pair: $\widehat{E}^{decoy,sequence}(i,j) = \sum_{a',b'} f(a')f(b')\, E^{decoy}(i,a',j,b')$, and $\sigma^{decoy,sequence}(i,j)$ is defined analogously to Eqs. (S3)–(S4) with the sum running over pairs $(a',b')$.

#### Configurational

For each native contact pair (i, j) only the direct pairwise energy is considered, with no background neighbours, i.e., $E^{native}(i,j) = E^{water}(i,j) + E^{elec}(i,j) + E^{burial}(i) + E^{burial}(j)$ (Eq. (S5) without the sums over $k$), where $i$ and $j$ are the residues involved in a contact (d<$r_c$=9.5). If the electrostatic term is disabled, $E^{\text{elec}}$ will be zero.

We sample 100,000 contact decoys by independently drawing a random distance *($r_{rand}$)* from the set of distances between all contacts, two residue densities ($\rho_1^{\text{rand}}$, $\rho_2^{\text{rand}}$, and two amino acid types from the protein's observed distribution ($\text{AA}_1^{\text{rand}}$, $\text{AA}_2^{\text{rand}}$), all drawn independently from different residues. Backbone positions of the native pair are not used. Note that the number of decoys to evaluate is a parameter that can be changed. It is by default 100,000 but if marginal speed-ups are needed in very large datasets, this number can be lowered to 2,000 without much cost in accuracy.

$$\begin{aligned} E^{\text{decoy}}(i,j) = {} & E^{\text{water}}(r_{\text{rand}}, \rho_1^{\text{rand}}, \rho_2^{\text{rand}}, \text{AA}_1^{\text{rand}}, \text{AA}_2^{\text{rand}}) \\ & + E^{\text{elec}}(\text{AA}_1^{\text{rand}}, \text{AA}_2^{\text{rand}}, r_{\text{rand}}) \\ & + E^{\text{burial}}(\text{AA}_1^{\text{rand}}, \rho_1^{\text{rand}}) + E^{\text{burial}}(\text{AA}_2^{\text{rand}}, \rho_2^{\text{rand}}) \end{aligned} \tag{S7}$$

where $i$ and $j$ are the residues involved in a contact (d<$r_c$=9.5). $r_{rand}$ is sampled from the distribution of all allowable pairwise distances in the protein.

#### Contact Pair Selection

For mutational and configurational modes, an amino acid pair (i, j) is included as a contact if:

(i) the $C_\beta$–$C_\beta$ distance (or C$\alpha$ for glycine) is less than 9.5 Å;

(ii) the sequence separation $|i - j| \geq 3$ (default value is 3; *--min-seq-sep* accepts any integer $> 0$, where 12 is often chosen to be compatible with early versions of the tool, but it is not recommended to change this value);

(iii) and both amino acid types pass the sequence mask (HETATM and UNK residues are excluded).

Only the upper triangle ($i < j$) is stored; the full symmetric contact matrix is recovered during energy evaluation. Note that residue density uses a *different*, stricter filter of $|i - j| > 1$ to exclude local backbone contacts from the burial density count (see Residue Density in Supplementary Methods) (Parra et al., 2016).

#### PDB fixing

Raw PDB files often contain non-standard amino acids, HETATM records, and missing heavy atoms. We apply the following preprocessing to match FrustratometeR conventions:

Non-standard amino acid residues were mapped to their corresponding canonical amino acid types prior to frustration calculations, including selenomethionine (MSE) to methionine (MET), protonation-state variants of histidine (HID, HIE, HIP, HSD, HSE, HSP) to histidine (HIS), disulfide-bonded, sulfenic-acid, and deprotonated cysteine variants (CYX, CY1, CYM) to cysteine (CYS), carbamylated and neutral lysine variants (KCX, LYN) to lysine (LYS), and protonated glutamate and aspartate (GLH, ASH) to glutamate (GLU) and aspartate (ASP), respectively. Remaining HETATM records and UNK residues are excluded (Parra et al., 2016).

Alternate conformations. Alternate conformations are resolved by selecting the first occupancy conformer.

Missing C$\beta$ atoms. Missing C$\beta$ atoms are placed using CHARMM36 (Huang and MacKerell, 2013) ideal internal coordinates from the backbone N, C$\alpha$, and C atoms, matching the MODELLER (Webb and Sali, 2016) placement used in FrustratometeR: $\vec{D} = \vec{C}_\alpha + \ell \cos\theta\, \hat{bc} + \ell \sin\theta \cos\phi\, \hat{y} - \ell \sin\theta \sin\phi\, \hat{n}$, with bond length $\ell = 1.5220$ Å, $\angle$C–C$\alpha$–C$\beta$ = 111.09°, dihedral N–C–C$\alpha$–C$\beta$ = 123.23°. Glycine is exempt since it has no C$\beta$ and uses C$\alpha$ as its centroid throughout.

Missing backbone oxygen atoms. Missing carbonyl O atoms are placed using ideal peptide-bond geometry from C$\alpha$, C, and the next residue's N.

#### GPU Implementation and Memory Management

All three frustration modes are formulated as batched PyTorch tensor operations over a batch dimension B (number of proteins), so all decoy energies are computed in a single forward pass without any Python-level loop over decoys.

We use PyTorch Lightning (Falcon et al., 2020) and a PyTorch Dataset data structure to load structures and handle batching on the CPU, while energy evaluation runs on one or more GPUs. In all modes, per-protein wall time is dominated by structure loading and feature computation rather than GPU energy evaluation.

Variable-length proteins are zero padded using a boolean mask. The pairwise C$\beta$–C$\beta$ (or C$\alpha$ for glycine) distance matrix is initially computed as a dense [B, L, L] float32 tensor; however, only the contact pairs within a given cutoff (9.5 Å default) are stored for energy function calculations. For proteins with L > 300, contacts are found using a GPU-accelerated fixed-radius spatial search in torch-cluster v1.6.3 (Fey et al., 2025; Fey and Lenssen, 2019). Since mutational mode is much more computationally expensive, we also chunk the background water energy, background electrostatic field, and decoy energy calculations to avoid out of memory errors, without changing the results.

#### AWSEM energy terms

The AWSEM hamiltonian depends on different factors that are explained below:

#### Burial Energy

The burial energy measures how deep each amino acid is buried using three overlapping density windows:

$$E^{burial}(i, a) = -\frac{k^{burial}}{2} \sum_{s=1}^{3} \gamma_{a,s}^{burial}\, \theta_s(\rho_i) \tag{S8}$$

Where $i$ is the residue index, $a$ is the residue type, and $\gamma_{a,s}^{\mathrm{burial}}$ is the burial statistical potential for amino acid type a in shell s, and the well function for each shell is:

$$\theta_s(\rho) = \tanh\left[\kappa(\rho - \rho_s^{\min})\right] + \tanh\left[\kappa(\rho_s^{\max} - \rho)\right] \tag{S9}$$

with $k^{\text{burial}} = 1.0$, $\kappa = 4.0$, $\rho_s^{\min} = [0, 3, 6]$, $\rho_s^{\max} = [3, 6, 9]$ (neighbour count units) (Davtyan et al., 2012; Papoian et al., 2004).

#### Water Energy

Each contact pair (i, j) has both water-mediated and protein-mediated interactions across two distance wells. The local density of both amino acids determines if a contact is water-exposed:

$$\sigma^{\text{water}}(i,j) = \frac{1}{4}\left(1 - \tanh[\eta_\sigma(\rho_i - \rho_0)]\right) \times \left(1 - \tanh[\eta_\sigma(\rho_j - \rho_0)]\right) \tag{S10}$$

$$\sigma^{\text{protein}}(i,j) = 1 - \sigma^{\text{water}}(i,j) \tag{S11}$$

The well function for each distance range is:

$$\theta^{\text{well}}(r; r_{\min}, r_{\max}) = \frac{1}{4}\left(1 + \tanh[\eta(r - r_{\min})]\right) \times \left(1 + \tanh[\eta(r_{\max} - r)]\right) \tag{S12}$$

The total water energy is:

$$\begin{aligned} E^{\text{water}}(i,j,r) = -\Big[ & \gamma^{\text{direct}}_{a_i,a_j}\,\theta^{\text{well}}(r;\, r^{\text{direct}}_{\min}, r^{\text{direct}}_{\max}) \\ & + \Big(\big(\sigma^{\text{protein}}(i,j)\,\gamma^{\text{protein}}_{a_i,a_j} + \sigma^{\text{water}}(i,j)\,\gamma^{\text{water}}_{a_i,a_j}\big) \\ & \quad \times\, \theta^{\text{well}}(r;\, r^{\text{mediated}}_{\min}, r^{\text{mediated}}_{\max})\Big)\Big] \end{aligned} \tag{S13}$$

where $\gamma$ tables are statistical potentials taken directly from AWSEM (Davtyan et al., 2012; Papoian et al., 2004). Parameters: $\eta_\sigma = 7.0$, $\rho_0 = 2.6$, $\eta = 5.0$, $r^{\text{direct}} = [4.5, 6.5]$ Å, $r^{\text{mediated}} = [6.5, 9.5]$ Å. r can either be the true distance, rij, between residues or i or j, or in the case for the configurational decoys, a random distance drawn from the distribution of allowable distances of the protein.

#### Electrostatics Energy

Charged amino acid pairs interact via a screened Coulomb potential (Debye–Hückel):

$$E^{\text{elec}}(i,j,r) = k_{ij}\frac{q_i q_j}{r}e^{-r/\lambda_D} \tag{S14}$$

where $q$ is the amino acid charge (D, E: $-1$; K, R: $+1$; all others: 0), $\lambda_D = 10.0$ Å is the Debye screening length, and $k_{ij} = 4.15$ kcal·mol$^{-1}$·Å. This is applied to all charged pairs with |i−j| $\geq 2$ regardless of distance; uncharged pairs contribute zero (Parra et al., 2016). $r$ can either be the true distance, $r_{ij}$, between residues or $i$ or $j$, or in the case for the configurational decoys, a random distance drawn from the distribution of allowable distances of the protein.

##### Residue Density

Residue density measures how many neighbours each amino acid has within a distance well, and is used to determine whether a contact is water-mediated or protein-mediated:

$$v_\theta(d) = \frac{1}{4}\left(1 + \tanh[\eta(d - r_{\min})]\right) \times \left(1 + \tanh[\eta(r_{\max} - d)]\right) \tag{S15}$$

$$\rho_i = \sum_{\ell:\,|\ell - i| > burial\_min\_sep} \theta^{\text{well}}(d_{i\ell};\, r_{\min}, r_{\max}) \tag{S16}$$

where $\theta^{\text{well}}$ is the same as defined above (see Water Energy) with $burial\_min\_sep = 1$, $r_{\min} = 4.5$ Å, $r_{\max} = 6.5$ Å. The centroid is $C_\beta$ for all amino acids except glycine, which uses $C_\alpha$ (Papoian et al., 2004).
